# *Anopheles* bionomic, insecticide resistance and malaria transmission in southwest Burkina Faso: a pre-intervention study

**DOI:** 10.1101/638551

**Authors:** Dieudonné Diloma Soma, Barnabas Muhugnon Zogo, Anthony Somé, Bertin N’Cho Tchiekoi, Domonbabele François de Sales Hien, Hermann Sié Pooda, Sanata Coulibaly, Jacques Edounou Gnambani, Ali Ouari, Karine Mouline, Amal Dahounto, Georges Anicet Ouédraogo, Florence Fournet, Alphonsine Amanan Koffi, Cédric Pennetier, Nicolas Moiroux, Roch Kounbobr Dabiré

**Author notes:** These authors contributed equally to this work (Anthony Somé, Bertin N’Cho Tchiekoi, Domonbabele François de Sales Hien, Hermann Sié Pooda). Corresponding author (DDS).

## Abstract

**Background:** The present study presents results of entomological surveys conducted to address the malaria vectors bionomic, insecticide resistance and transmission prior to the implementation of new strategies complement long-lasting insecticidal nets (LLINs) in the framework of a randomized control trial in southwest Burkina Faso.

**Methods:** We conducted entomological surveys in 27 villages during the dry cold season (January 2017), dry hot season (March 2017) and rainy season (June 2017). We carried out hourly catches (from 17:00 to 09:00) inside and outside 4 houses in each village using the Human Landing Catch technique. Mosquitoes were identified using morphological taxonomic keys. Specimens belonging to the *Anopheles gambiae* complex and *Funestus* Group were identified using molecular techniques as well as detection of *Plasmodium falciparum* infection and insecticide resistance target-site mutations.

**Results:** Eight *Anopheles* species were detected in the area. *Anopheles funestus s.s* was the main vector during the dry cold season. It was replaced by *Anopheles coluzzii* during the dry hot season whereas *An. coluzzii* and *An. gambiae s.s.* were the dominant species during the rainy season. Species composition of the *Anopheles* population varied significantly among surveys. All researched target site mutation of insecticide resistance (*kdr-w, kdr-e* and *ace-1*) were detected in all members of the *An. gambiae* complex of the area but at different frequencies. We observed early and late biting phenotypes in the main malaria vector species. Entomological inoculation rates were 0.087, 0.089 and 0.375 infected bites per human per night during dry cold season, dry hot season and rainy season, respectively.

**Conclusion:** The intensity of malaria transmission was high despite the universal coverage with LLINs. We detected early and late biting phenotypes in the main malaria vector species as well as physiological insecticide resistance mechanisms. These vectors might mediate residual transmission. These data highlight the need to develop complementary tools in addition to LLINs in order to better control resistant malaria vectors and to monitor insecticide resistance.

## Background

The World Health Organization (WHO) has reported 219 million malaria cases and 435 000 deaths worldwide in 2017 [1]. Significant progress has been made in the era of malaria control and elimination between 2000 and 2017 in all regions of the world [1]. The number of malaria cases worldwide has decreased by 62% between 2000 and 2015 but seems to rebound since 2016 [1–3]. Sub-Saharan Africa accounted for about 92% of cases and deaths in 2017 [1]. In 2017, the health ministry of Burkina Faso recorded 11.9 million cases and 4144 deaths attributed to malaria [4].

Malaria control is mainly based on symptomatic and preventive treatments (with artemisinin-based combination therapies: ACTs) and vector control. Vector control aims at reducing malaria transmission by targeting the *Anopheles* mosquitoes that transmit *Plasmodium* parasites. Vector control relies mostly on the mass distribution of long-lasting insecticidal nets (LLINs) [3]. It was estimated that LLINs have contributed to 68 % of the decline in malaria cases observed between 2000 and 2015 [5] despite moderate use rates [6]. However, the emergence of physiological [7–9] and behavioral [10,11] insecticide resistance mechanisms in *Anopheles* mosquitoes, as observed in most parts of Africa, could compromise the effectiveness of LLINs and explains the recent rebound in malaria cases.

Therefore, it urges to increase or complement the protection provided by the LLINs with complementary strategies. Complementary strategies exists but before being included into strategic plans by national malaria control programs (NMCPs), supported by international donors and implemented in endemic countries, they need to be evaluated through rigorous and independent process [12]. In this context, Institut de Recherche en Sciences de la Santé (IRSS), Institut de Recherche pour le Développement (IRD) and Institut Pierre Richet (IPR) have been funded to conduct the REACT project to evaluate in Burkina Faso and Côte d’Ivoire four complementary strategies to LLINs. These strategies included: i) larviciding with *Bacillus thuringiensis israelensis* (Bti) to target immature stages of *Anopheles* species, Indoor residual spraying (IRS) with pirimiphos-methy to target endophilic malaria vectors, information, education, community (IEC) to improve LLINs use iv) ivermectin administration to animals, a One health approach to tackle zoophagic behavior of malaria vectors and to improve animal health of malaria vectors and to improve animal health. This study presents the result of entomological surveys conducted to address the malaria vectors bionomic, insecticide resistance and transmission prior to the implementation of complementary strategies to LLINs in the framework of a randomized control trial (RCT) in southwest Burkina Faso.

## Materials and methods

### Study site and design

The study was conducted in the Diébougou health district located in southwest Burkina Faso (**Fig 1**). The natural vegetation is mainly wooded savannah dotted with clear forest gallery. The climate is tropical with two seasons: one dry season from October to April and one rainy season from May to September. The average annual rainfall is about 1200 mm. The dry season is “cold” from December to February (with average minimal and maximal temperatures of 18°C and 35°C, respectively) and “hot” from March to April (with average minimal and maximal temperatures of 25°C and 40°C, respectively). Agriculture (cotton growing and cereals) is the main economic activity in the area, followed by artisanal gold mining and production of coal and wood [13,14]. The study took place in 27 villages that were selected considering accessibility during the rainy season, a size of 200-500 inhabitants and a distance between two villages higher than two kilometers.

**Fig 1.**
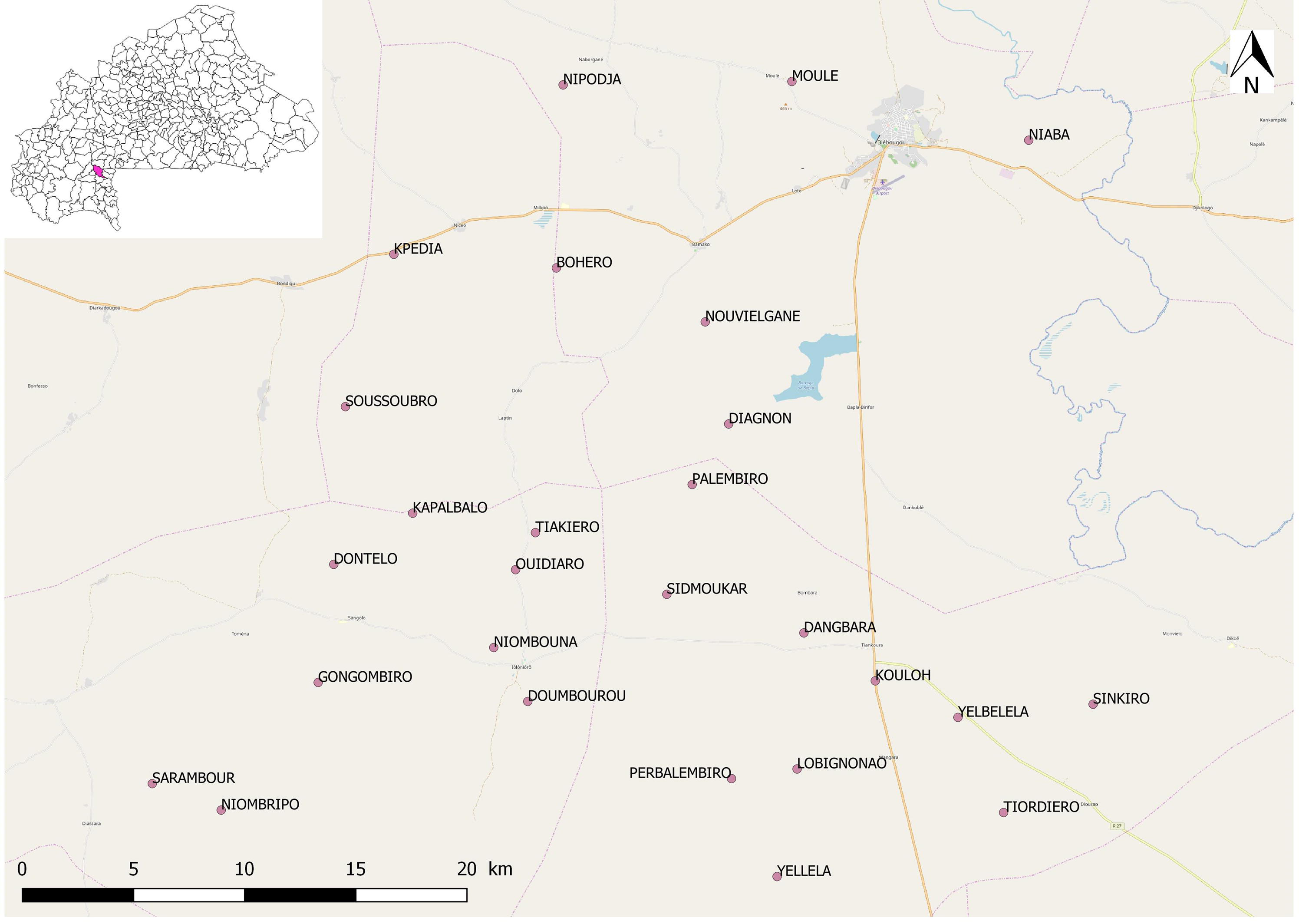
Study area and mosquito collections sites

### Mosquito collection and determination

We carried out three surveys of mosquito collection in January 2017 (dry cold season), March 2017 (dry hot season) and June 2017 (rainy season) using Human Landing Collections (HLC). Mosquitoes were collected from 17:00 to 09:00 both indoors and outdoors at 4 sites per villages for one night per survey. Due to a long duration of collection (16h), two teams of 8 collectors worked in each village and shifted at 01:00. Collectors were rotated among the collection points every hour. The distance between collection sites was at least 50 m. The distance between indoors and outdoors collection points in one site was at least 10 m. Independent staff supervised rotation of the mosquito collection and every time they checked quality.

### Morphological identification and dissection

Mosquitoes were morphologically identified on the field to genus and species levels using morphological keys [15,16]. A subsample of 100 non blood-fed *Anopheles spp.* individuals was randomly selected per survey and per village to determine their parous status [17]. All females belonging to the *Anopheles* genus were stored in individual tubes with silicagel and preserved at -20°C for further analyses.

### Molecular analysis

DNA extracted from head-thorax of *Anopheles spp.* individuals was used to detect *Plasmodium falciparum* infection using quantitative polymerase chain reaction (PCR) assay [18]. Individuals belonging to the *Anopheles gambiae* complex and the *Funestus* Group were identified to species by PCR [19–21].

PCR assay were carried out on all mosquitoes belonging to the *An. gambiae* complex to detect the L1014F (*kdr-w*) [22], the L1014S (*kdr-e*) [23] and the G119S (*ace-1*) [24] mutations. These mutations confer insecticide resistance to the pyrethroids for the *kdr-w* and *kdr-e* and to carbamates and organophosphates for the *ace-1*.

### Parameters measured

We calculated the human biting rate (HBR; the number of vectors’ bites per human per night (b.h^-1^.n^-1^), the *Plasmodium* infection rates (PIR: the proportion of *Anopheles* infected by *P. falciparum*), the entomological inoculation rate (EIR; the number of infected bites per human per night (ib.h^-1^.n^-1^ that is the product of the HBR with the PIR), the parous rate (PR; the proportion of parous females over the total dissected) and the endophagy rate (ER; the proportion of *Anopheles* females collected indoors) for each *Anopheles* species and for overall *Anopheles spp.*

### Statistical analyses

We assessed HBR, ER, PIR, EIR and PR using generalized mixed effect models (GLMM) with collection points and villages as nested random intercept. All models were fitted using the ‘glmer’ function of the package ‘lme4’ [25] run in the R software [26]. For all models, we performed backward stepwise deletion of the fixed terms followed by Likelihood ratio tests. Term removals that significantly reduced explanatory power (*p* < 0.05) were retained in the minimal adequate model. We used the post-hoc Tukey method to do multiple comparison among modalities of the fixed terms using the ‘emmeans’ function of the ‘emmeans’ package [27]. We considered main effects and interactions of the fixed terms in all the models.

We performed HBR and EIR analyses using negative-binomial models fitted on nightly counts of all *Anopheles* and of *Anopheles* individuals found to be infected by *P. falciparum*, respectively. Fixed terms for these counts models were the survey and the collection position (indoor or outdoor).

We performed PIR, ER and PR analyses using binomial models fitted on individual status of the *Anopheles* (infected *vs*. susceptible, collected indoors *vs.* outdoors or parous *vs*. nulliparous, respectively). Fixed terms for these binomial models were the survey and the individual species.

The relationship between *Anopheles* species composition and surveys was tested using a Pearson’s Chi-squared test of independence with simulated p-value.

We assessed nightly activity of each major *Anopheles* species by comparing their Median Catching Time (MCT), which represents the time at which 50% of the individuals were collected [10], using a Kruskall-Wallis test and a Dunn’s post-hoc test for multiple comparisons (function ‘dunnTest’ of the ‘VIA’ package) in R [28].

We compared the genoptypes frequencies of *kdr* and *ace-1* mutations among surveys and species using the G-test [29] implemented in Genepop 4.7 and run in R [30]. Genotypic frequencies at the *kdr* and *ace-1* mutations were tested for conformity to Hardy-Weinberg equilibrium using the “Exact HW test” [31]. In case of disequilibrium, we tested heterozygote excess and deficiency using the score test [32].

### Ethics approval and consent to participate

The protocol of this study was reviewed and approved by the Institutional Ethics Committee of the Institut de Recherche en Sciences de la Santé (IEC-IRSS) and registered as N°A06/2016/CEIRES. Mosquito collectors and supervisors gave their written informed consent. They received a vaccine against yellow fever as a prophylactic measure. Collectors were treated free of charge for malaria according to WHO recommendations.

## Results

## *Anopheles* densities and composition

During the three surveys, we collected a total of 2591 mosquitoes belonging to four genus: *Anopheles spp.* (n=1936, 74.72%), *Aedes spp.* (n=481, 18.56%), *Culex spp.* (n=161, 6.21%) and *Mansonia spp.* (n=13, 0.50%) (**Table 1**). Among the 1530 *Anopheles* individuals morphologically identified as members of the *An. gambiae* complex and proceeded by PCR, 1131 were *An. coluzzii*, 325 were *An. gambiae s.s.* and 74 were *An. arabiensis. Anopheles* morphologically identified as members of the *Funestus* Group and successfully proceeded by PCR were all (n=250) *An. funestus s.s*. Other *Anopheles* species found during our surveys were *Anopheles nili* (n=3), *Anopheles pharoensis* (n=23), *Anopheles rufipes* (n=1) and *Anopheles squamosus* (n=1) (**Figs 2A-C**). We successfully identified by PCR 92.84% (1530/1648) of the *An. gambiae s.l*. individuals and 96.15% (250/260) of the members of the *Funestus* Group. The average HBR of *Anopheles spp.* mosquitoes was 1.30 and 1.34 b.h^-1^.n^-1^ (bites per human per night) during surveys one and two (dry season, RR_1-2_ = 1.15 [0.78, 1.71], p=0.67), respectively, that was significantly lower than during survey three in rainy season (6.31 b.h^-1^.n^-1^; RR_1-3_= 0.090 [0.063, 0.130], Tukey’s p< 0.0001; RR_2-3_= 0.078 [0.054, 0.114], Tukey’s p< 0.0001) (**Fig 3**).

**Table 1.** Abundance and diversity of mosquito species in the Diébougou area during pre-intervention surveys

| Mosquitoes species | Dry cold season | Dry hot season | Rainy season | Total |
| --- | --- | --- | --- | --- |
| *Aedes aegypti* | 2 | 1 | 103 | 106 |
| *Aedes africanus* | 0 | 0 | 6 | 6 |
| *Aedes fowleri* | 0 | 0 | 100 | 100 |
| *Aedes furcifer* | 0 | 0 | 101 | 101 |
| *Aedes luteocephalus* | 0 | 0 | 16 | 16 |
| *Aedes opok* | 0 | 0 | 16 | 16 |
| *Aedes vexans* | 0 | 0 | 89 | 89 |
| *Aedes vittatus* | 0 | 6 | 41 | 47 |
| *Anopheles gambiae s.l.* | 35 | 264 | 1349 | 1648 |
| *Funestus* Group | 230 | 25 | 5 | 261 |
| *Anopheles nili* | 1 | 0 | 2 | 3 |
| *Anopheles pharoensis* | 14 | 1 | 8 | 23 |
| *Anopheles rufipes* | 1 | 0 | 0 | 1 |
| *Anopheles squamosus* | 0 | 0 | 1 | 1 |
| *Culex cinereus* | 1 | 9 | 15 | 25 |
| *Culex decens* | 0 | 1 | 2 | 3 |
| *Culex poicilipes* | 0 | 1 | 1 | 2 |
| *Culex quinquefasciatus* | 21 | 42 | 62 | 125 |
| *Culex univittatus* | 1 | 0 | 5 | 6 |
| *Mansonia africana* | 4 | 4 | 0 | 8 |
| *Mansonia uniformis* | 2 | 0 | 3 | 5 |
| Total | 312 | 354 | 1925 | 2591 |

Bars indicate 95% confidence intervals; Red dots show the mean densities per survey; Survey 1: dry cold season, Survey 2: dry hot season, Survey 3: rainy season.

During the dry cold season (survey 1), *An. funestus s.s* was the most abundant species representing 78.64% (n=221/281), followed by *An. gambiae s.s* 6.04%, (17/281), *An. coluzzii* 4.98%, (14/281) and *An. pharoensis* 4.98%, (14/281) (Fig 2A). The other species were found at very low percentages (∼1%) (**Fig 2A**). The relative abundance and species composition of the *Anopheles* population varied from one village to another. We collected *Anopheles* mosquitoes in 19 villages out of the 27 surveyed with the higher densities registered in Diagnon (n= 149, 4 species) and Kpédia (n = 39, 3 species) (**Fig 2A**). We observed the higher *Anopheles* species diversity in Niaba (n=15) where 6 species were identified. At the opposite in several villages (n= 9), all the *Anopheles* found were either *An. funestus s.s, An. coluzzii* or *An. pharoensis*.

**Fig 2A.**
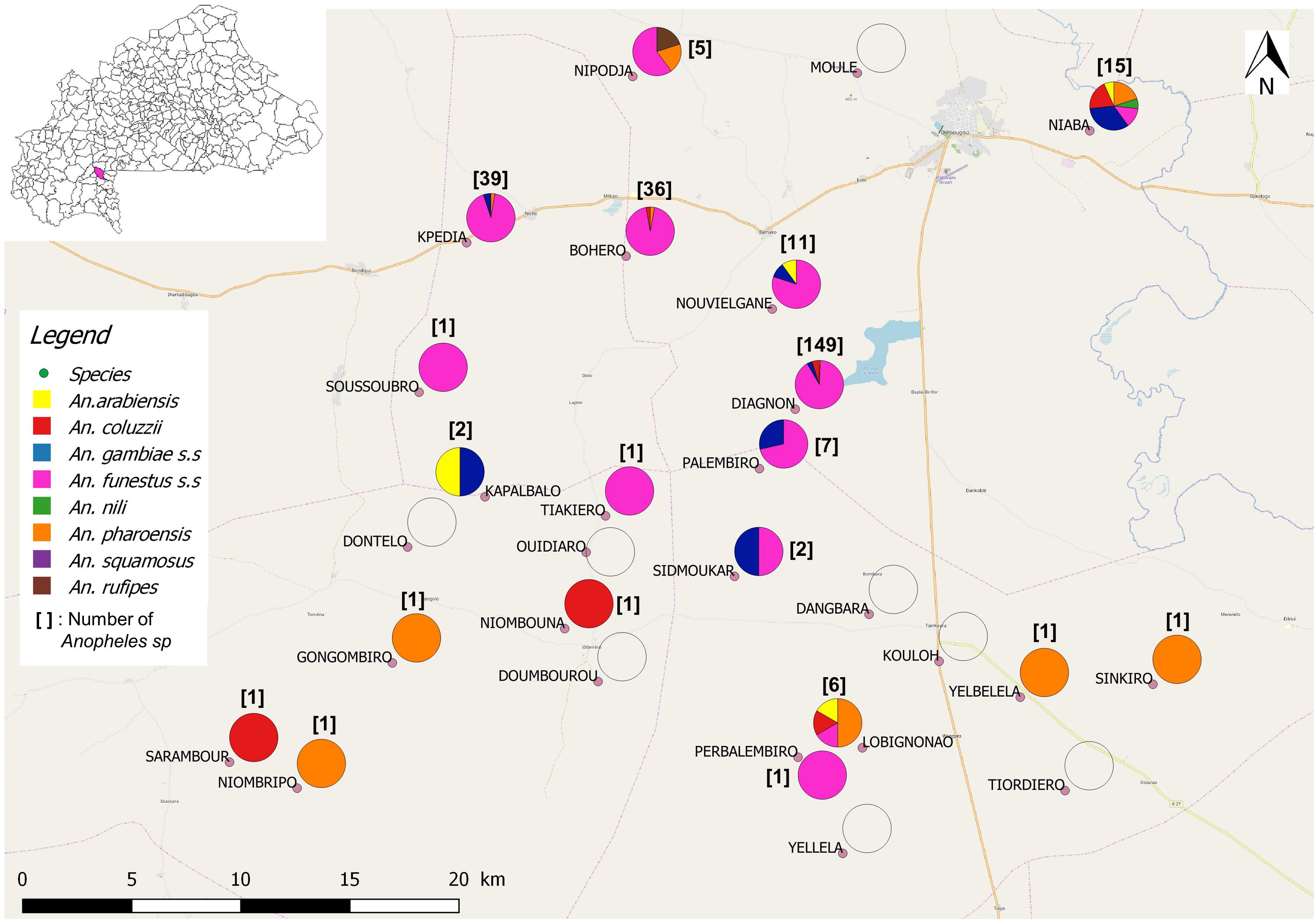
Map of *Anopheles* densities and composition during dry cold season

**Fig 2B.**
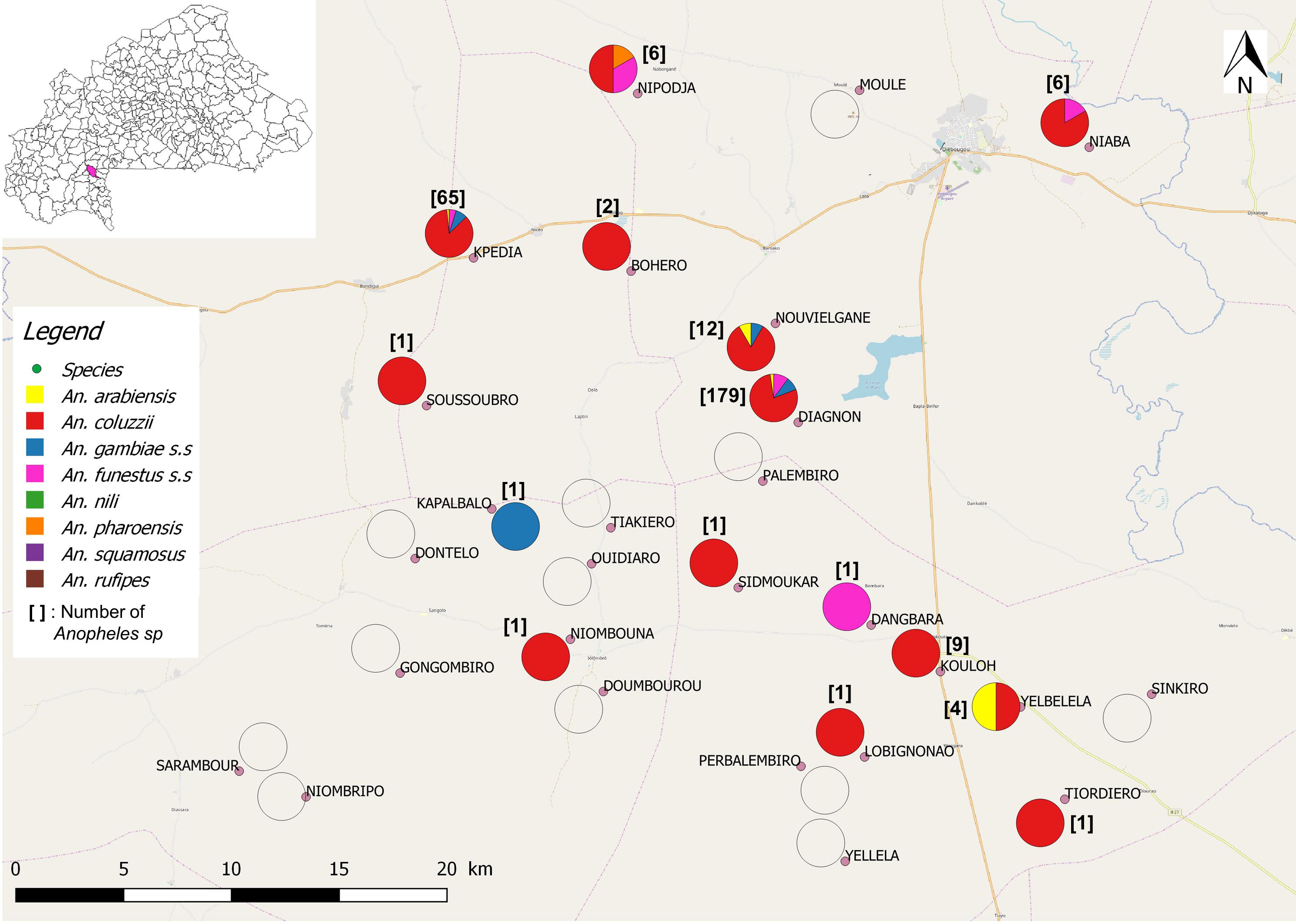
Map of *Anopheles* densities and composition during dry hot season

**Fig 2C.**
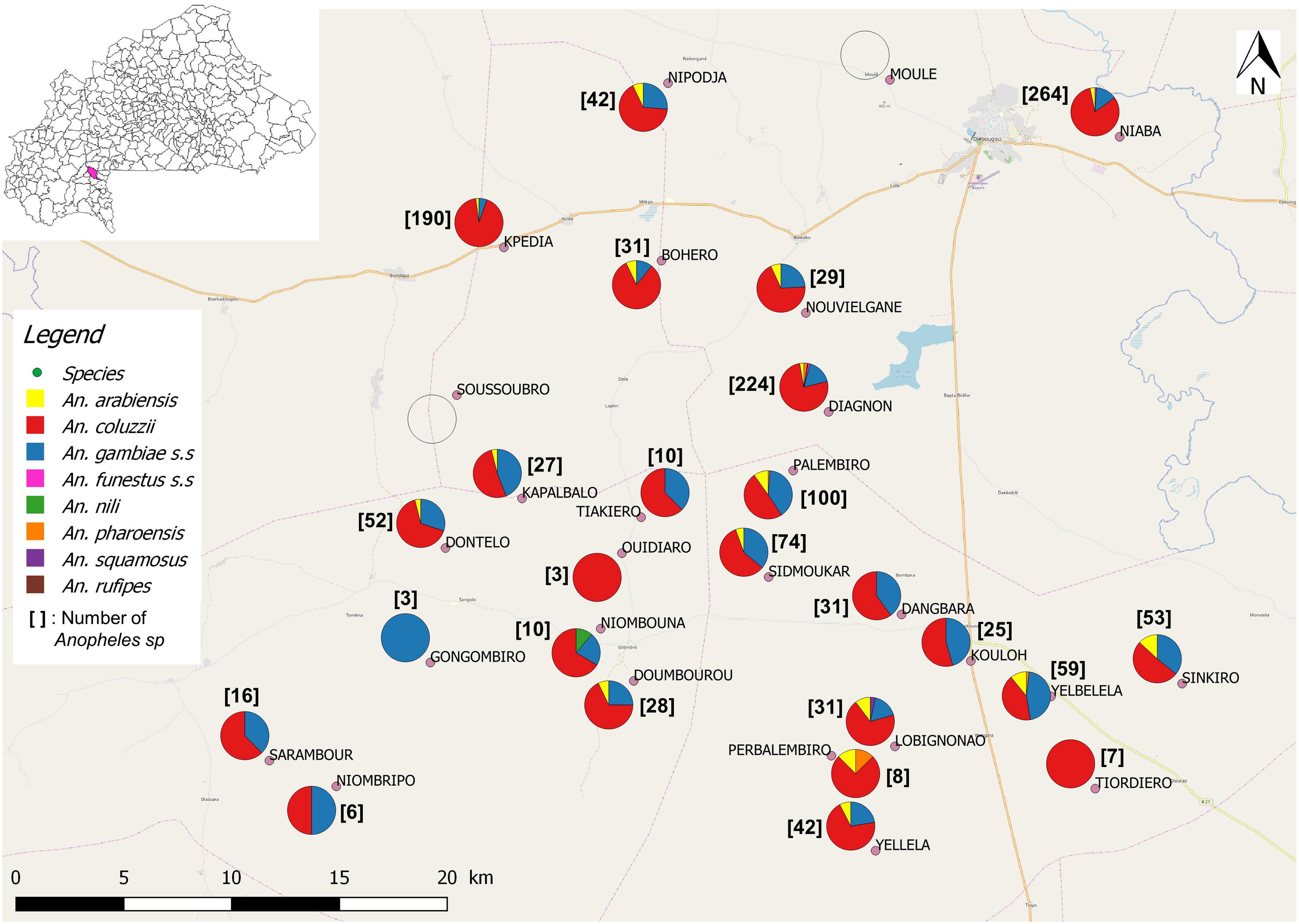
Map of *Anopheles* densities and composition during rainy season

**Fig 3.**
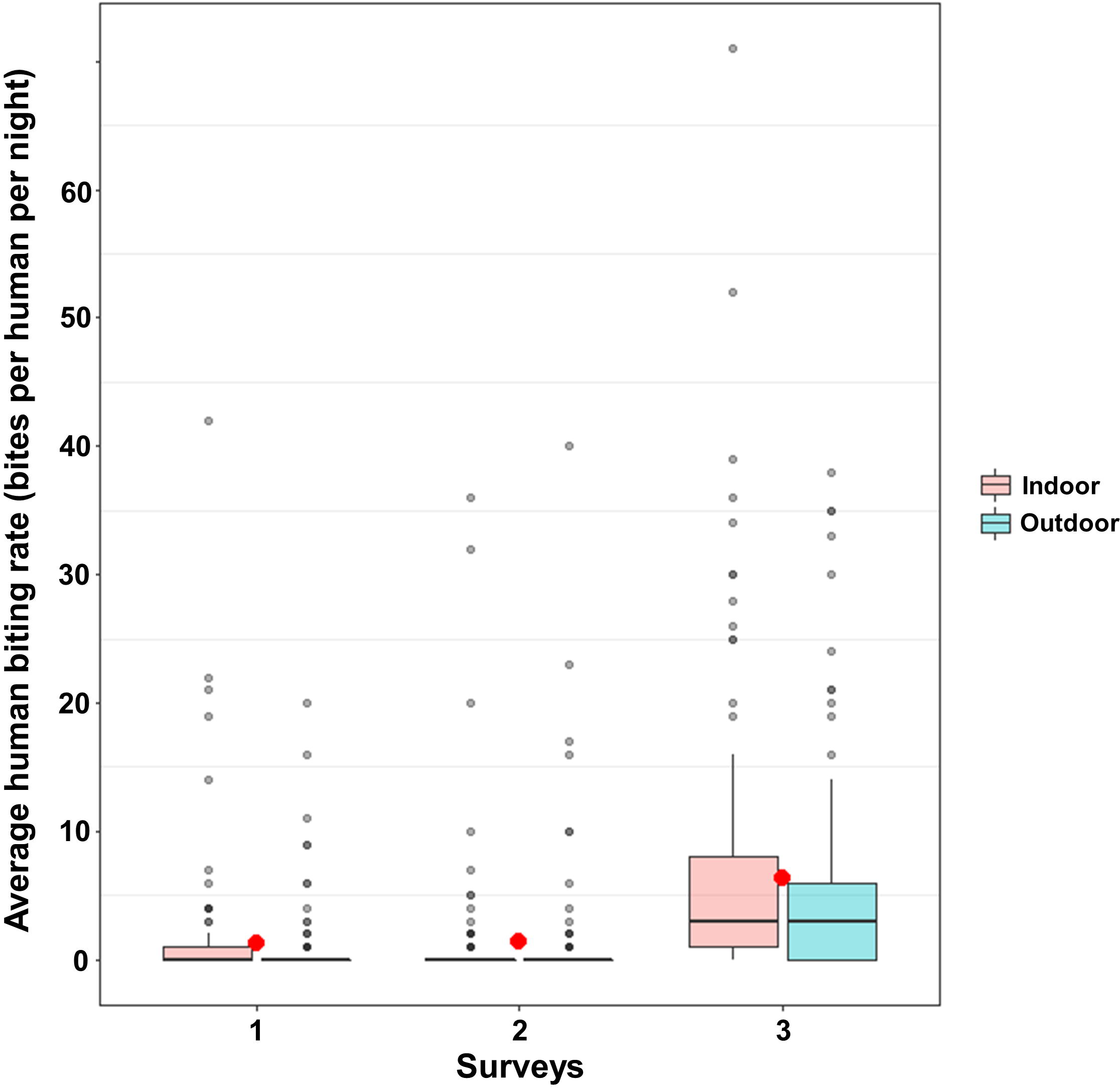
Average human biting rate of *Anopheles* mosquitoes collected outdoors and indoors. During survey 2 in the dry hot season, *An. coluzzii* (75.86%, n=220/290) almost totally replaced *An. funestus s.s* (8.27%, n=24/290) while proportions of other species do not vary substantially. During this survey, *Anopheles* mosquitoes were collected in only 15 villages (**Fig 2B**). Higher densities and species diversities were observed in Diagnon (n = 179, 4 species) and Kpédia (n = 65, 4 species) (**Fig 2B**). We identified 3 *Anopheles* species in both Nouvielgane and Nipodja villages and 2 *Anopheles* species in Yelbelela and Niaba. In all other villages, only one *Anopheles* species was present at low densities.

During survey 3 in the rainy season, *An. coluzzii* remained the major species (65.71%, n=897/1365) followed by *An. gambiae s.s* (20.95%, n=286/1365) and *An. arabiensis* (4.54%, n=62/1365). During this survey, *An. funestus s.s* fall under 1% of the total. We collected *Anopheles spp.* mosquitoes in 25 villages out of the 27 surveyed (**Fig 2C**). The higher densities were observed in Niaba (n = 264, 4 species), Diagnon (n = 224, 6 species) and Kpédia (n=190, 3 species). We identified 4 *Anopheles* species in Lobignonao (n=31), Palembiro (n=100), Yelbelela (n=59), 2 *Anopheles* species in Dangbara (n=31), Kouloh (n=25), Niombripo (n=6), Sarambour (n=16), Tiakiero (n=10), and 1 *Anopheles* species in Gongombiro (n=3), Ouidiaro (n=3), Tiordiero (n=7). In the remaining villages, 3 *Anopheles* species was sampled. No *Anopheles spp* mosquitoes were collected in Sousoubro and Moulé (**Fig 2C**). Species composition of the *Anopheles* population varied significantly among surveys (Pearson’s Chi-squared test, X^2^=1339.7, simulated p-value = 0.0005).

### Mosquito biting behavior

Overall, endophagy rate (ER) of *Anopheles spp.* [IC95%] was 63.23% [57.50-68.96], 50.18% [44.27-56.09] and 57.18% [54.44-59.90] during the dry cold, dry hot and rainy seasons, respectively (**Table 2**).

**Table 2.** *Anopheles* species composition and densities

| Species | Dry cold season |  | Dry hot season |  | Rainy season |  |
| --- | --- | --- | --- | --- | --- | --- |
|  | Indoor | Outdoor | Indoor | Outdoor | Indoor | Outdoor |
| *An. arabiensis* | 2 | 2 | 2 | 6 | 31 | 31 |
| *An. coluzzii* | 5 | 9 | 106 | 114 | 525 | 372 |
| *An. gambiae s.s* | 14 | 3 | 16 | 6 | 159 | 127 |
| *An. funestus s.s* | 146 | 75 | 14 | 10 | 3 | 2 |
| *An. nili* | 1 | - | - | - | 1 | 1 |
| *An. pharoensis* | 3 | 11 | - | 1 | 2 | 6 |
| *An. rufipes* | 1 | - | - | - | - | - |
| *An. squamosus* | - | - | - | - | - | 1 |
| Total | 170 | 98 | 136 | 131 | 690 | 509 |
| % Indoors [95% CI] | 63.23 [57.50-68.96] |  | 50.18 [44.27-56.09] |  | 57.17 [54.44-59.90] |  |

During survey 1 (dry cold season), *An. funestus s.s* (marginal mean ER [95% CI] = 64.8% [56.6, 72.2], p= 0.0005) and *An. gambiae s.s.* (marginal mean ER = 83.7% [59.0, 94.9], p = 0.012) were significantly endophilic. During this survey, ER of other species were not different than 50% (p>0.25).

During survey 2 (dry hot season), ERs of *An. funestus s.s* and *An. gambiae s.s* decreased to 56.2% [35.5, 75.0] and 71.1% [48.3, 86.6], respectively, no longer different than 0.5 (p = 0.56 and p=0.069, respectively). During this survey, ER of other species were also not different than 50% (p>0.14).

During survey 3 (rainy season), the ER of *An. coluzzii* increased significantly (RR_2-3_ = 0.61 [0.41, 0.89], p = 0.0066) to 59.3% [54.7, 63.8] significantly higher than 50% (p=0.0001). During this survey, ER of other species were also not different than 50% (p>0.12).

The median catching times of *An. coluzzii, An. gambiae s.s*, and *An. pharoensis* were recorded between 01:00 and 02:00 while those of *An. funestus s.s* and *An. arabiensis* were one hour later (between 02:00 and 03:00; **Fig 4**). These differences were significant between *An. funestus s.s* and both *An. coluzzii* and *An. gambiae s.s* (Dunn’s multiple comparison test p-values = 0.01 and 0.004, respectively). *An. coluzzii, An. gambiae s.s, An. pharoensis* and *An. funestus s.s* showed early biting activity (beginning at 18:00) but *An. arabiensis* (beginning at 21:00). A late biting activity (after 06:00) was observed with *An. coluzzii, An. gambiae s.s* and *An. funestus s.s* (**Fig 4**).

**Fig 4.**
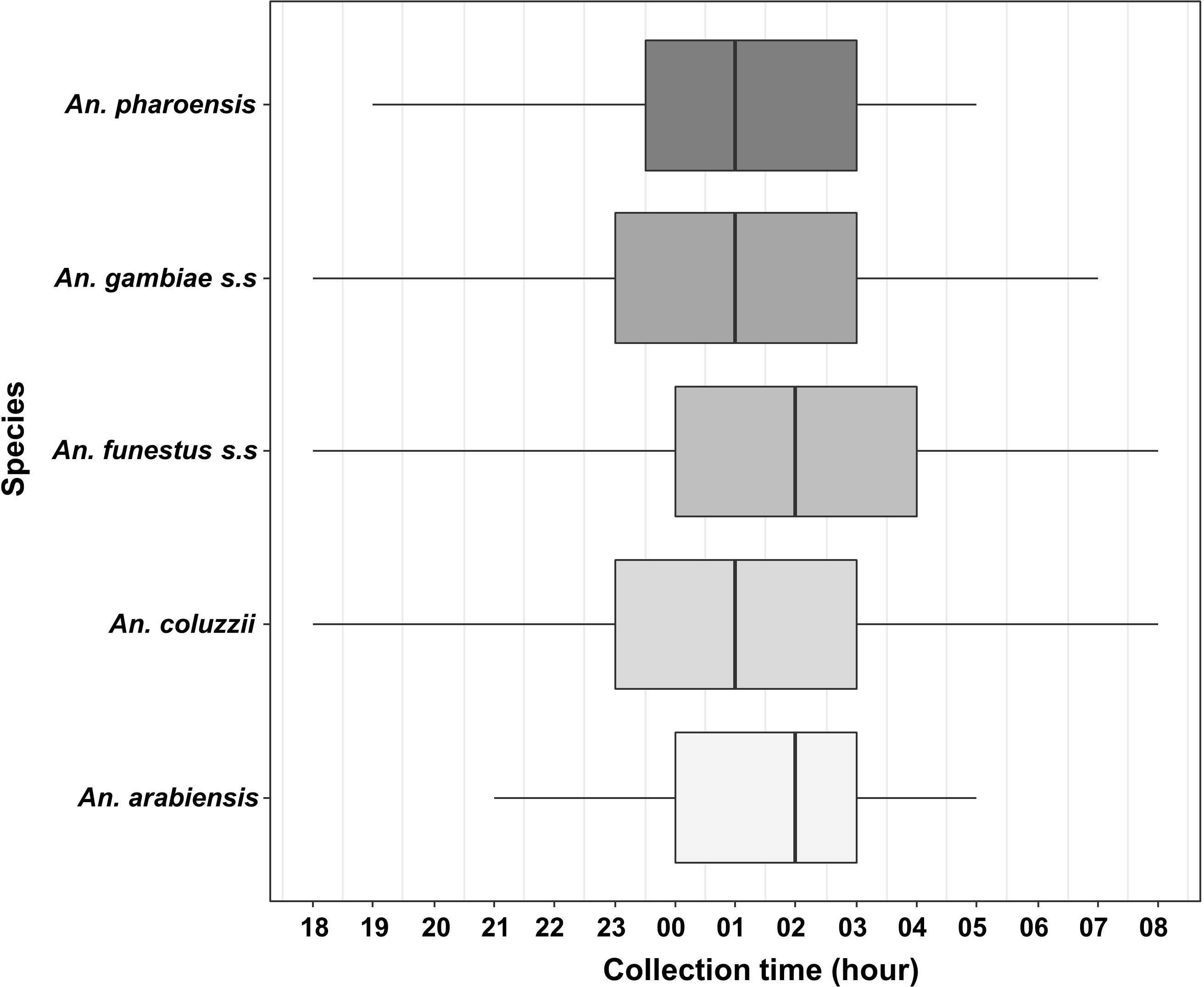
Median catching time of *Anopheles*. Boxes indicate 1st-3rd quartile and median hours of biting activity. Whiskers indicate 2.5–97.5 percentiles

## *Plasmodium* infection and transmission intensity

We analyzed, a total of 1808 *Anopheles* for the research of *P. falciparum* infection. The overall *Plasmodium* infection rate was 6.98 % [2.03-14.38] (19/272), 6.90 % [1.03-11.35] (19/275) and 6.42 % [0.00-6.49] (81/1261) in dry cold season, dry hot season and rainy season, respectively. None *An. nili, An. pharoensis, An. rufipes* and *An. squamosus* individuals tested were infected. The PIR did not vary significantly among surveys (Tukey’s tests, p-values > 0.09). The PIR of *An. gambiae s.s* was lower than that of *An. funestus s.s* (OR=0.20 [0.05, 0.90], p= 0.03). No other differences between species have been evidenced (p-values > 0.23).

Entomological inoculation rate (EIR) was 0.375 [0.30-0.44] infected bites per human per night (ib.h^-1^.n^-1^) during survey 3 (rainy season) significantly higher than 0.087 ib.h^-1^.n^-1^ [0.06-0.10] measured during survey 1 (dry cold season; RR_1-3_=0.25 [0.11, 0.56], p=0.0002) and

0.089 ib.h^-1^.n^-1^ [0.05-0.11] measured during survey 2 (dry hot season; RR_2-3_=0.23 [0.10, 0.51], p=0.0001).

**Table 3.** Entomological transmission parameters

| species | Dry cold season |  | Dry hot season |  | Rainy season |  |
| --- | --- | --- | --- | --- | --- | --- |
|  | PIR (%) | EIR | PIR (%) | EIR | PIR (%) | EIR |
| *An. arabiensis* | 0.00 | 0.000 | 12.50 | 0.005 | 1.61 | 0.005 |
| *An. coluzzii* | 14.29 | 0.009 | 7.73 | 0.079 | 6.35 | 0.264 |
| *An. gambiae s.s* | 11.76 | 0.009 | 4.55 | 0.005 | 8.04 | 0.106 |
| *An. funestus s.s* | 6.79 | 0.069 | 0.00 | 0.000 | 0.00 | 0.000 |
| *An. nili* | 0.00 | 0.000 | - | - | 0.00 | 0.000 |
| *An. pharoensis* | 0.00 | 0.000 | 0.00 | 0.000 | 0.00 | 0.000 |
| *An. rufipes* | 0.00 | 0.000 | - | - | - | - |
| *An. squamosus* | - | - | - | - | 0.00 | 0.000 |
| **Total** | **6.98** | **0.087** | **6.90** | **0.089** | **6.42** | **0.375** |
| **[95% CI]** | **[2.03-14.38]** | **[0.06-0.10]** | **[1.03-11.35]** | **[0.05-0.11]** | **[0.00-6.49]** | **[0.30-0.44]** |

PIR: *Plasmodium* infection rate; EIR: entomological inoculation rate; CI: Confidence interval

## Physiological age

We dissected 966 *Anopheles* for determination of parous rate. *Anopheles* parous rate was 76.00% [68.51-83.48], 78.80 [72.28-85.32] and 66.66% [63.14-70.18] in the dry cold season, dry hot season and rainy season respectively (**S1 Table**). The average parous rate of *Anopheles* mosquitoes during surveys two (dry hot season) was significantly higher than during survey three (rainy season) (OR=2.1, [1.21-3.70], p-value=0.0051). Overall, parous rate was 76.38%, 68.97%, 67.24%, 72.65%, 77.77% and 100% for *An. funestus s.s, An. arabiensis, An. coluzzii, An. gambiae s.s, An. pharoensis* and *An. nili* respectively (**S1 Table**). The parous rate did not differ significantly among the species (χ ^2^ = 2.2, df = 3, p=0.51).

## Frequencies of L1014F *kdr*, L1014S *kdr* and G119S *ace-1* mutations in *An. gambiae* s.l

Numbers of individuals of each genotype of the three mutations and their frequencies in *An. arabiensis, An. gambiae s.s.* and *An. coluzzii* are presented for each survey (**Table 4**).

**Table 4.** Allele frequency of *kdr* L1014F, *kdr* L1014S and *ace-1* G119S mutations in *Anopheles gambiae* s.l. populations

| Species | Period | Genotypes *kdr-w* |  |  |  |  |  | Genotypes *kdr-e* |  |  |  |  |  | Genotypes *ace-1* |  |  |  |  |  |
| --- | --- | --- | --- | --- | --- | --- | --- | --- | --- | --- | --- | --- | --- | --- | --- | --- | --- | --- | --- |
|  |  | N | SS | RS | RR | f(1014F) | p(HW) | N | SS | RS | RR | f(1014S) | p(HW) | N | SS | RS | RR | f(119S) | p(HW) |
| *An. arabiensis* | Dry cold season | 4 | 0 | 4 | 0 | 0.500<sup>a</sup> | 0.3141 | 4 | 4 | 0 | 0 | 0.000<sup>a</sup> | - | 3 | 3 | 0 | 0 | 0.000<sup>a</sup> | - |
|  | Dry hot season | 8 | 5 | 3 | 0 | 0.188<sup>a</sup> | 1.0000 | 8 | 8 | 0 | 0 | 0.000<sup>a</sup> | - | 8 | 6 | 2 | 0 | 0.125<sup>b</sup> | 1.0000 |
|  | Rainy season | 56 | 35 | 18 | 3 | 0.214<sup>a</sup> | 0.6978 | 53 | 22 | 31 | 0 | 0.292<sup>b</sup> | 0.0022 | 61 | 61 | 0 | 0 | 0.000<sup>a</sup> | - |
| *An. coluzzii* | Dry cold season | 14 | 1 | 9 | 4 | 0.607<sup>ab</sup> | 0.3164 | 14 | 14 | 0 | 0 | 0.000<sup>a</sup> | - | 14 | 12 | 2 | 0 | 0.071<sup>a</sup> | 1.0000 |
|  | Dry hot season | 219 | 37 | 63 | 119 | 0.687<sup>a</sup> | 0.0000 | 218 | 217 | 1 | 0 | 0.002<sup>a</sup> | - | 219 | 211 | 8 | 0 | 0.018<sup>a</sup> | 1.0000 |
|  | Rainy season | 823 | 176 | 389 | 258 | 0.550<sup>b</sup> | 0.1938 | 743 | 453 | 289 | 1 | 0.196<sup>b</sup> | 0.0000 | 888 | 849 | 39 | 0 | 0.022<sup>a</sup> | 1.0000 |
| *An. gambiae* s.s | Dry cold season | 17 | 0 | 1 | 16 | 0.971<sup>a</sup> | - | 17 | 17 | 0 | 0 | 0.000<sup>a</sup> | - | 16 | 9 | 7 | 0 | 0.219<sup>a</sup> | 0.5446 |
|  | Dry hot season | 22 | 8 | 2 | 12 | 0.591<sup>b</sup> | 0.0000 | 22 | 22 | 0 | 0 | 0.000<sup>a</sup> | - | 22 | 17 | 5 | 0 | 0.114<sup>a</sup> | 1.0000 |
|  | Rainy season | 258 | 16 | 26 | 216 | 0.888<sup>a</sup> | 0.0000 | 226 | 213 | 13 | 0 | 0.031<sup>a</sup> | 1.0000 | 282 | 179 | 97 | 6 | 0.193<sup>a</sup> | 0.1253 |

N: number of mosquitoes; SS: homozygous susceptible; RS: heterozygous; RR: homozygous resistant; f(1014F): frequency of the 1014F resistant *kdr* allele; f(1014S): frequency of the 1014S resistant *kdr* allele; f(119S): frequency of the 119S resistant *ace-1* allele ; p(HW) : exact Hardy-Weinberg test p-value; '-': not determined; In each species and mutation, allelic frequencies carrying the same superscript letter do not differ significantly (G-test $p > 0.05$ ).

In *An. arabiensis*, the *kdr-w* mutation did not vary significantly among surveys (exact G test p-values > 0.15) nor among villages (exact G test p-values > 0.08). The population did not differ significantly from the HWE (exact HW test p > 0.31) whatever the survey.

The *kdr-w* frequency in *An. coluzzii* during the dry cold season was 0.61, it increased (not significantly, exact G test p = 0.41) to 0.69 during dry hot season and then decreased significantly to 0.55 during the rainy season (exact G test p<0.001). During the dry hot season, when the frequency of the *kdr-w* mutation was the highest, the population was not at the HW equilibrium (exact HW test p< 0.001) due to heterozygote deficiency (p<0.001) observed in the village of Diagnon where most of the individuals (131/220) where collected.

In *An. gambiae s.s*, the *kdr-w* mutation frequency was 0.97 during dry cold season and significantly decrease in dry hot season to 0.59 (exact G test p < 0.001). During the rainy season when densities were very high, the *kdr-w* frequency rose up to 0.88. That was significantly higher than during dry hot season (exact G test p < 0.001) but not different than during dry cold season (p = 0.12). This population was not at the HWE for the *kdr-w* mutation (exact HW test p < 0.001) due to heterozygote deficiency (p < 0.001). Heterozygote deficiency was observed in most of the villages during each survey (**S1 File**).

In *An. arabiensis*, the *kdr-e* mutation was detected only during the rainy season at a frequency of 0.30. The frequency of the *kdr-e* mutation did not vary significantly among villages (exact G test p > 0.10). A significant deviation of HWE was observed (exact HW test p = 0.002) due to heterozygote excess in rainy season (p= 0.001). However, we did not observe this heterozygote excess at the village scale (**S2 File**).

In *An. coluzzii*, the *kdr-e* mutation was not detected during the dry-cold season and only one heterozygous individual was collected during the dry hot season corresponding to a frequency of 0.002. The frequency increased significantly to 0.12 during the rainy season (exact G test p < 0.001). A significant deviation of HWE was observed for the *kdr-e* mutation (exact HW test p < 0.001) due to heterozygote excess in most of the villages during each survey (**S3 File**).

In *An. gambiae s.s*, the *kdr-e* mutation was detected only during the rainy season at a frequency of 0.03. The *kdr-e* mutation do not vary significantly among villages (exact G test p > 0.11). The population did not differ significantly from the HWE (exact HW test p > 0.05) in rainy season.

In *An. arabiensis*, the *ace-1* he *kdr-e* mutation was detected only during the dry hot season at a frequency of 0.12. The frequency of the *ace-1* mutation did not vary significantly among villages (exact G test p > 0.48). The population did not differ significantly from the HWE (exact HW test p > 0.05) in dry hot season.

The frequency of the *ace-1* allele in *An. coluzzii* was 0.07, 0.02 and 0.02 in dry cold, dry hot and rainy seasons, respectively. There were no significant difference in the *ace-1* allele frequency among surveys (exact G test p > 0.11) nor among villages (exact G test p > 0.13). The *ace-1* allele frequency in *An. coluzzii* did not differ from the HWE (exact HW test p > 0.05).

In *An. gambiae s.s*, the frequencies of the *ace-1* allele were 0.22, 0.11 and 0.19 in dry cold, dry hot and rainy seasons, respectively. The frequency did not vary significantly among surveys (exact G test p > 0.23) nor among villages (exact G test p > 0.07). The population did not differ significantly from the HWE (exact HW test p > 0.12) whatever the survey.

## Discussion

This study showed that the malaria vector species abundance in the Diébougou area varied significantly according to the season. *Anopheles funestus s.s* was the predominant vector during the dry cold season (January 2017) but most individuals were collected in one village (Diagnon) that is close to swamps depending on the dam of Bapla. *Anopheles funestus s.s* densities were divided by 10, two month later during the dry hot season (March 2017) and it almost disappeared in June 2017 during the rainy season. *Anopheles funestus s.s* is known to breed in large permanent or semi-permanent pools preferentially with emergent vegetation on its margins [33,34]. In Burkina Faso, two chromosomal forms of *An. funestus* have been described named Folonzo and Kiribina [35,36]. Folonzo form of *An. funestus* is associated with the presence of water reservoirs containing natural vegetation, such as swamps. The swamps near Diagnon are upstream from the Bapla dam and become dry at the end of the dry season. This may explain why *An. funestus s.s* almost disappeared during the dry hot season and until swamps becomes flooded again.

*Anopheles coluzzii* was shown at very low densities during the dry cold season but became the predominant malaria vector species from the dry hot season (March 2017) to the rainy season (June 2017). During the dry-hot season, most of individuals were collected in Diagnon, the same village where *An. funestus s.s* densities were simultaneously falling. In Burkina Faso, *An. coluzzii* is known to breed in permanent or semi-permanent breeding sites [37,38]. Its presence in Diagnon during the dry season is certainly linked to the dam. However, it is not clear why *An. coluzzii* was present in very low densities during January 2017 and became numerous two months later. We hypothesize that the reduction of the breeding sites favorable to *An. funestus s.s* may have increased *An. coluzzii* competitiveness against *An. funestus s.s* around the Bapla dam.

Densities of *An. gambiae s.s.* and *An. arabiensis* were low during both dry season’s surveys and increased substantially during the rainy season (in a larger extent for *An. gambiae s.s*.). This is consistent with the preference of both these species to breed in temporary rain-dependent pools and puddles [34,37,39].

These four species (*An. funestus s.s, An. coluzzii, An. gambiae s.s* and *An. arabiensis)* were responsible for all the *P. falciparum* transmission recorded in this study. *Plasmodium* infection rate did not vary among surveys but EIR increase from an average of about 3 infected bites per human per month in dry (cold and hot) season to more than 10 bites per human per month during the rainy season. As PIR was stable over the study, the increase in EIR in rainy season was mathematically due to the increase in vectors densities. According to our results, malaria transmission persists all over the year in the study area despite high spatial disparities. Indeed, during the dry cold and dry hot seasons, the spatial distribution of *An. funestus s.s.* and *An. coluzzii*, that were the main malaria vectors, was restrained to a small number of villages that concentrated malaria transmission. During the rainy season, the spatial distribution of malaria vector (with predominance of *An. coluzzii* and *An. gambiae s.s.*) was more homogeneous.

In addition to these well-known primary malaria vectors, we recorded the presence of four other *Anopheles* species in low densities (*An. nili, An. pharoensis, An. rufipes, An. squamosus*). While we were not able to detect *Plasmodium* parasites in the few individuals belonging to these species, they are all potential vectors of *Plasmodium* [34,40–44]. They are considered secondary vectors because of their behavior (exophagy and/or zoophagy) [42] but are nevertheless likely to maintain residual levels of transmission [45]. Such residual transmission by secondary vector species may increase as a result of the implementation of vector control measures which are designed to target primary vectors. Indeed, drastic reduction of the major vector’s densities may freeing up ecological niches for secondary vectors [46–49]. Therefore, secondary vectors deserve particular attention [50]. Their bionomics and competence for *P. falciparum* transmission remain to be investigated.

Regarding the behavior of the main malaria vector species, we observed that malaria vectors were slightly but mainly endophagic. This indicates that indoor vector control measures (such as LLINs and IRS) are expected to target a significant part of the vector population but probably insufficient to stop the transmission. The peak of aggressiveness of the main *Anopheles* species occurred during the second part of the night. This observation is consistent with the usual patterns of hourly biting aggressiveness of these species. This period corresponds to the moments of deep sleep of the human populations who are potentially protected by LLINs [39,51]. However, we observed early- and late-biting phenotypes in the main malaria vector species (*An. coluzzii, An. funestus s.s*. and *An. gambiae s.s*). These phenotypes might mediate residual transmission and we might expect to see them selected by the massive use of insecticide-based vector control tools at night such as LLINs [10,52]. It is therefore crucial to monitor malaria vector behavior in this area when implementing malaria vector control strategies in order to track the possible emergence of behavioral resistances.

This study showed high parity rates of malaria vectors (> 60%) in the study area, regardless of the season. The parous rate was significantly lower in the rainy season than in the cold dry season and the hot dry season, indicating that young females were more prevalent in the rainy season. The lower parous rate of *Anopheles* vectors during the rainy season may be due to the presence of more breeding habitats which yield more nulliparous mosquitoes in vector populations [53]. It is also possible that a higher usage of LLINs during the rainy season as a result of a high nuisance and low temperatures might result in smaller proportions of parous vectors [54]. The three main insecticide resistance mutation *kdr-w, kdr-e* and *ace-1* have been detected in the three species of the *An. gambiae* complex but at varying frequencies among species and surveys. In June 2017 during the rainy season (when the populations were most abundant), the *kdr-w* mutation frequency was very high in *An. gambiae s.s.* (almost fixed, f=0.88). Since the discovering of this mutation [22] in *An. gambiae s.s*, the *kdr-w* frequencies have always been high in this species in Southwest Burkina Faso [7,9,55] indicating that the mutation has a low cost for this species or that strong selection pressures occur, such vector control strategies implementation (universal distribution of LLINs is implemented in Burkina Faso since 2010 [56]) and massive use of pesticides in agriculture [57]. Indeed, the intensive use of insecticides in cotton cultivation would be a major factor driving the selection of pyrethroid-resistant specimens in Southwest Burkina Faso [9]. Most of the populations of *An. gambiae s.s.* from this study showed heterozygote deficiency for the *kdr-w* mutation that could indicate a Whalund effect resulting from the presence of several populations within each given village. This might be explained by the colonization of tempory breeding sites by different vector populations during the rainy season as evidence the annual reappearance of *An. gambiae s.s.* in the Sahel after the dry season [58]. To note that no disequilibrium was observed for the *kdr-e* and *ace-1* mutations in this species indicating that these populations, if they exists, resemble the local population in terms of genotypic composition for these loci.

During the rainy season, the *kdr-w* mutation frequency was lower in *An. coluzzii* (f= 0.55) and *An. arabiensis* (f=0.21) than in *An. gambiae s.s*. But in counterpart, both *An. coluzzii* (f=0.19) and *An. arabiensis* (f=0.29) showed higher prevalence of the *kdr-e* mutations than *An. gambiae s.s.* (f=0.03). In *An. coluzzii* and *An. arabiensis*, a heterozygote excess was observed for the *kdr-e* mutation. This disequilibrium may result from a heterozygote advantage as it has been observed in mating in *An. coluzzii* specimen from Southwest Burkina Faso carrying the *kdr-w* mutation [59]. The causes of this disequilibrium remain to be explored.

Both *kdr* mutations are well “implanted” in the vector population of our study area. These mutations provide physiological resistance to the pyrethroids insecticides, the only family of compounds authorized to impregnate LLINs. Although the impact of pyrethroids resistance on the efficacy of malaria control remains disputable [60], the development and the implementation of resistance management strategy need to be encouraged.

Regarding the *ace-1* mutation that confers resistance to carbamates and organophosphates insecticides, it was present in the three species at a moderate frequency in *An. gambiae s.s.* but at very low frequencies in *An. coluzzii* and *An. arabiensis*. In Burkina Faso, the three species may express high frequencies of the *ace-1* mutation depending on the location [61]. In our study area, *An. gambiae s.s.* seems to suffer more selection pressure possibly due to a wide use of carbamates and organophosphates particularly for cotton growing.

## Conclusions

Malaria transmission in the Diébougou area was mainly due to *An. funestus s.s, An. coluzzii, An. gambiae s.s* and *An. arabiensis* with high spatio-temporal heterogeneities. The intensity of malaria transmission was high despite the presence of LLINs. We observed early and late biting phenotypes in the main malaria vector species. These phenotypes might mediate residual transmission. Three mutations *kdr-w, kdr-e* and *ace-1* were present in the three species of the *An. gambiae* complex with high frequency variability between species. These data highlighted the need to complement malaria control interventions to establish efficient resistance management plan (integrated vector control approach) and monitor insecticide resistance (both physiological and behavioral).

## Acknowledgements

We thank all participants of the study, especially technicians at the IRSS-DRO for their technical assistance. We acknowledge the Burkina Faso Ministry of Health, particularly Dr. Dembélé Henri that facilitated the data collection. We are also, grateful to the villagers of all sites for their kind collaboration and hospitality.

## Supporting information

**S1 Table. Parous rate of *Anopheles***

**S1 File. Hardy Weinberg test when H1= heterozygote deficit in *An. gambiae s.s* populations**

**S2 File. Hardy Weinberg test when H1= heterozygote excess in *An. Arabiensis* populations**

**S3 File. Hardy Weinberg test when H1= heterozygote deficit in *An. coluzzii* populations**

### Data Availability Statement

The datasets used during this article are available at the Institut de Recherche en Sciences de la Santé, Bobo-Dioulasso, Burkina Faso and will be made easily available on request, when required.

### Funding

This work was part of the REACT project, funded by the French Initiative 5 % Expertise France (N° 15SANIN213).

