## Supplementary material for "*Anopheles* bionomic, insecticide resistance and malaria transmission in southwest Burkina Faso: a pre-intervention study": S1 Table

**S1 Table. Parous rate of *Anopheles***

| Species | Dry cold season | | |  | Dry hot season | | |  | Rainy season | | |
| --- | --- | --- | --- | --- | --- | --- | --- | --- | --- | --- | --- |
|  | Number dissected | parous | Parous rate (%) [95% CI] | | Number dissected | parous | Parous rate (%) |  | Number dissected | parous | Parous rate (%) |
| *An. arabiensis* | 1 | 0 | 0 |  | 2 | 1 | 50 [0.0- 100] |  | 26 | 19 | 73,07 [56.02-90.12] |
| *An. coluzzii* | 5 | 3 | 60 [17.05-100] |  | 115 | 93 | 80,87 [73.68-88.05] |  | 457 | 292 | 63,89 [59.49-68.29] |
| *An. gambiae s.s* | 9 | 8 | 88,89 [68.35-100] |  | 17 | 11 | 64,7 [41.98-87.42] |  | 197 | 143 | 72,58 [66.35-78.81] |
| *An. funestus s.s* | 108 | 83 | 76,85 [68.89-84.80] |  | 16 | 13 | 81,25 [62.12-100] |  | 3 | 1 | 33,33 [0.0-86.67] |
| *An. nili* | - | - | - |  | - | - | - |  | 1 | 1 | 100 |
| *An. pharoensis* | 2 | 1 | 50 [0-100] |  | 1 | 1 | 100 |  | 6 | 4 | 66,66 [28.94-100] |
| Total | 125 | 95 | 76 [68.51-83.48] |  | 151 | 119 | 78,8 [72.28-85.32] |  | 690 | 460 | 66,66 [63.14-70.18] |
